# Active EB1 surges promote tubulin influx into the growing outer segments of the bipartite olfactory cilia in *Drosophila*

**DOI:** 10.1101/2024.09.10.612170

**Authors:** Riddhi Girdhar Agarwal, Saishree Iyer, Ayan Barbora, Yogesh Gadgil, Swadhin Jana, Krishanu Ray

## Abstract

Like a photoreceptor cilium, the sensory cilia have a complex bipartite architecture containing 9+0 connecting cilium at the base and a singlet microtubule-supported, highly membranous outer segment, essential for the receptor display. How such diverse cilia morphology and underlying microtubule cytoskeleton develops remains unclear. Here we show that individual olfactory cilium, inside the large basiconic sensilla in developing *Drosophila* antenna, grows in episodic steps following several pulsatile influxes of tubulin. Each tubulin influx event is preceded by transient elevations of a microtubule-stabilising protein, the End-binding protein 1 (EB1). Additionally, EB1 is found to specifically interact with the tail domain of *Drosophila* KLP68D, an orthologue of the kinesin-2β motor subunit, *in vitro*. Finally, the loss of EB1 in olfactory neurons preceding the growth surges reduces the tubulin influx as well as arrests the olfactory cilia assembly and stability. These findings suggest a novel mechanism of bipartite cilia assembly.

## Introduction

The primary cilium is a microtubule-based structure that displays a variety of receptors on the ciliary membrane (1,2). It extends from the cell surface like an antenna, perceives environmental cues, and modulates intracellular signalling pathways. While most cilia appear as long cylindrical extensions supported by typical (9+0 and 9+2) arrangements of doublet microtubules, called axoneme, certain sensory cilia have ciliary membranes organised in structures such as discs of the rod photoreceptor outer segment (3), the branched arboreal arrangement with singlet microtubules in the ciliary outer segment (OS) of the winged cilia of *C. elegans* (4), and the long-branched extension of the olfactory cilia of *Drosophila* (5,6). These modifications increase the surface area to volume ratio to accommodate more receptors and enhance the sensory reception of the environmental cues. All these cilia have a bipartite cytoskeletal organisation consisting of a connecting cilium (CC) supported by the 9+0 arrangement of doublet microtubules at the base and singlet microtubule-bearing distal/outer segment (OS). Although the growth patterns of the canonical, 9+0 axoneme-bearing primary cilium are extensively studied (7–10), there are only a handful of reports on bipartite cilia growth. Therefore, how the elaborate ciliary outer segments grow is still unclear.

The bipartite olfactory cilia, expressed by Olfactory Sensory Neurons (OSNs) that innervate *sensilla basiconica* in the *Drosophila* antenna (henceforth referred to as the basiconic cilia) serve as an excellent model system for studying complex cilia assembly in a cell-type-specific manner. These cilia assemble over a prolonged period (nearly sixty hours) during pupal development (5), offering a valuable window to monitor protein localizations in different parts of the growing cilium in vivo. A previous study has defined the patterns of the length and volume growth of ciliary OS, indicating active recruitment of structural proteins in response to developmental cues, during the assembly of the ciliary OS (5). Several studies also highlight the role of kinesin-2 motor in trafficking proteins inside cilia, transporting different types of cargo from the ciliary base to the tip (5,8,11–13). Kinesin-2 is likely to interact with the αTubulin1A/D isoforms directly through its tail domain (13) and with the βTubulin subunit through the IFT74/81 complex (10). IFT74 and IFT81 are crucial for the growth and stability of motile and primary cilia in mammals (14–16). Also, mutations in the kinesin-2 motor subunit gene, *Klp64D*, halt progressive tubulin enrichment in developing olfactory cilia (13) and reduce the number of singlet microtubule-bearing OS branches (5,13). Together these results suggested that active tubulin transport could control ciliary growth characteristics and function. However, how these transports are regulated remains one of the key unresolved issues.

A primary cilium grows by incorporating tubulin dimers at the tip of the axoneme (17), which remains relatively stable throughout the resting phase and disassembles at the beginning of cell division (18,19). This stability is achieved through the decoration of several Microtubule Associated Proteins (MAPs) along the axoneme (17). One such MAP, the End-binding protein 1 (EB1), known to stabilize the plus end-directed growth of microtubules in cell culture (20), is found to decorate the singlet microtubule at the distal ends of the cilia expressed on the MDCK cells (21). EB1 organizes the +TIP complex by recruiting structurally and functionally diverse partner proteins onto the growing plus ends of microtubules (20). This interaction promotes the persistent growth of microtubules by decreasing the catastrophe frequency (22,23). Loss of EB1 disrupts formations of primary cilia in the human fibroblast cells bearing the 9+0 axoneme (24) and the *Chlamydomonas* flagella bearing the 9+2 axoneme (25). The protein is enriched at the growing tips of the flagella (25) and the base of the primary cilia in a resting phase cell (26). Therefore, it is construed that EB1 entry could stabilise the plus-end-directed growth of the ciliary microtubule in the same manner as indicated in the *in vitro* studies and promote further movement of tubulin towards the growing tip.

Therefore, to understand the mechanism of the bipartite olfactory cilia assembly and the role of kinesin-2 in the process, we studied the patterns of tubulin and EB1 entry into two different types of bipartite cilia innervating the *s. basiconica*. The results indicate that the ciliary OS grows with periodic surges of EB1 followed by that of tubulin, indicating an active recruitment and movement of these proteins through the cilium during the OS growth. Loss of EB1 dampened the tubulin surges and disrupted the stability of the ciliary OS. EB1 is found to interact with the tail domain of the KLP68D (kinesin-2β) subunit of *Drosophila* kinesin-2 in vitro. Further, tissue-specific Klp68D RNAi during the growth phases abrogated the EB1 surges and tubulin enrichment in the ciliary OS and severely affected their morphology. Our results suggest that developmentally regulated active recruitment of ciliary assembly proteins assists the growth of different regions of complex cilia. Further, EB1 appeared to play a distinct role in the tubulin entry process, quite different from the estimated plus end-tracking function of the protein.

## Results

### The outer segments of olfactory cilia inside the large s. basiconica grow in episodic phases

Two to four OSNs innervate each large *s. basiconica* (Fig. 1A and A’) and all of them expressed the *chaGal4* driver. Each of these olfactory cilium bears a bipartite microtubule cytoskeleton. The connecting cilium at the base is supported by a 9+0 arrangement of microtubule doublets, called axoneme. Whereas both the inner (IS) and the outer (OS) segments are supported by multiple singlet microtubules (Fig 1B-C’) (12). A previous study reported that these olfactory cilia assembled in an episodic manner during 36-96 hours after pupa formation (hrs APF) with a significant volume growth marked by the accumulation of GFPαTubulin84B, expressed using *chaGal4* driver, between 84 and 90 hrs APF (Fig S1A-S1B). We further noticed that this late growth phase consists of two separate spurts at 85-86 and 89-90 hrs APF (Fig S1B) and it reconciled with similar episodic enrichments of Jupiter, a microtubule-associated protein, in the OS during this period (Fig S1 C), indicating that the volume growth of the OS is likely to be supported by microtubule assembly.

**Fig 1.**
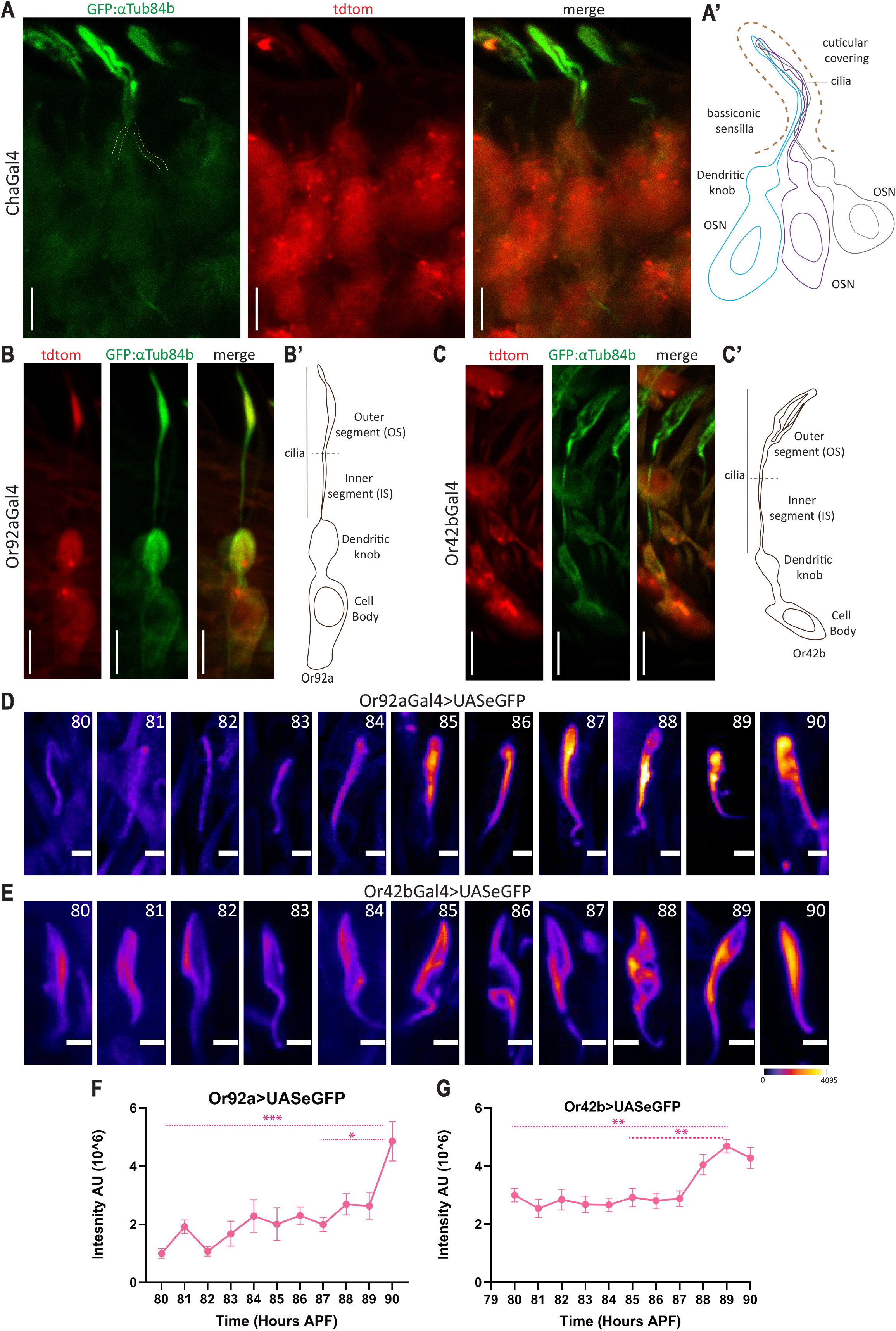
Growth profiles of sensory cilia expressed on the Or92A and Or42B olfactory sensory neurons. **A)** Expression of the GFPTub84B and soluble tdTomato in the Olfactory Sensory Neurons (OSN) in the chaGal4 UAS-GFPαTub84B; UAS-tdTom background marks the cilia (green) and the OSN cell bodies innervating the ab1-type s.bassiconica as indicated by the tracing (A’). **B and C)** Images of a typical Or92A **(B)** and a typical Or42B **(C)** OSNs, each marked by the GFPαTub84B (green) and tdTomato (red) expressions due to the Or92AGal4 and Or42BGal4 drivers, respectively, depict the morphological distinctions of cilia expressed on these two types of OSNs innervating the ab1-type s. basiconica. Scale bars indicate 5 µm and the tracings **(B’ and C’)** indicate different parts of each neuron. **D and E)** Developmental changes in the morphologies of Or92A and Or42B cilia, marked by the neuron-specific expression of eGFP in Or92A and Or42B OSNs, respectively. The intensity changes at different developmental stages after pupa formation (APF) are highlighted using a false colour heat map (FIRE, ImageJ®) and the scale bars indicate 2 µm in each panel. **F and G)** Integrated fluorescence intensity (mean +/-S.E.M) of eGFP in Or92A and Or42B cilia during the 80-90hr APF window. The pairwise significance of difference was estimated using Kruskal-Wallis ANOVA test, p-values (*p<0.05, **p<0.01, and ***p<0.001) are shown on the plots. Note a distinctly different growth pattern of each type of cilia.

The growth of highly branched singlet microtubules in the ciliary OS would require rapid and profuse microtubule assembly. EB1, a plus end-binding protein that stabilises the microtubule growth by reducing the catastrophe frequency in vitro (22) and decorating the singlet microtubules in the primary cilia expressed on the MDCK cells (21), could potentially harness this rapid microtubule growth. We found that the EB1GFP marked the entire OS of the olfactory cilia and its levels increased in an episodic manner during the 84-90 hrs APF (Fig S1D). Surprisingly, unlike GFPαTubulin84B and JupiterGFP, the EB1GFP levels peaked an hour before each of the microtubule growth phases and attenuated during the growth phases (Fig S1D).

The distinctive coincidence of GFPαTubulin84B influx, which is expressed using an ectopic *chaGal4* promoter and endogenously expressed JupiterGFP, ruled out the possibility of an overexpression artefact. Also, the EB1GFP, expressed using the same *chaGal4* promoter entered the cilia much before the GFPαTubulin84B. Hence, these observations suggested that episodic tubulin and EB1 entries in segregated phases may promote microtubule growth within the cilia. Since each large *s. basiconica* is innervated with multiple cilia, and due to the limitations in the optical resolution, it was unclear whether each cilium grows in synchronised phases or independently of the others. Further, this observation highlighted a peculiar pulsatile EB1 movement into and out of the cilia.

### Olfactory cilia extended from the Or92A and Or42B neurons, grow independently of each other

To investigate how cilia from individual neurons innervating the basiconic sensilla grow, we expressed eGFP in either the Or92a or Or42b neurons (Fig 1B and C). Although both cilia have a bipartite structure, the Or42B ciliary OS appears much broader with two principal branches whereas the Or92A ciliary OS has a bulbous shape. Ectopic eGFP expression in the Or92A (*Or92AGal4>UAS-eGFP*) and Or42B OSNs (*Or42BGal4>UAS-eGFP*) revealed distinct growth patterns (Fig 1D, E). The volume of Or42b ciliary OS increased gradually until 89 hrs APF and then nearly doubled at 90 hrs APF (Fig 1F). In contrast, the volume of the Or92a ciliary OS increased significantly (∼2x) during 87-89 hrs APF (Fig 1G). These observations suggested that each olfactory cilium within the same *s. basiconica* shaft grows independently, following unique patterns. This observation also suggested that collectively the Or42B and Or92A cilia are likely contribute to the second episodic growth of composite cilia observed during 89-90 hrs APF.

### Loss of EB1 disrupts cilia stability and microtubule growth inside the ciliary outer segment

Next, we sought to address the role of pulsatile EB1 entry and exit in the cilia growth during the growth of OS. EB1 family members consist of small (28-30 kDa) microtubule-associated proteins that were shown to preferentially associate with the growing ends of the microtubule filaments during interphase and metaphase (27,28) and regulate the plus-end-directed growth by stabilizing the nascent microtubule protofilaments (22). EB1 family members form obligate homo/heterodimers and bind to the target proteins through the consensus SxIP motif (29,30), which stabilizes the +TIP complex and microtubule growth (28,31). EB1 is enriched at the distal ends of the Chlamydomonas flagella (25) and the tips of the motile cilia of bronchial epithelial cells (24). Although EB1 was not detected in the primary cilia of epithelial cells, loss of EB1 eliminated the primary cilia and flagella (24,25). Altogether, the evidence suggested that EB1 would be essential for the cilia and flagella formation. However, how it assists in cilia formation remained unclear.

Like the EB1GFP, the EB1tdTomato, expressed in the OSNs by *chaGal4*, labeled the entire OS inside the *s. basiconica* (Fig S2Aa). No EB1tdTomato was present along the inner segment (IS) and in the CC region. The lengthwise EB1 decoration of the ciliary OS is consistent with a recent observation, which indicated decoration along the singlet microtubules at the distal end of the cilia expressed on the MDCKII cell line (21). In comparison, the dominant-negative version of EB1 (EB1^DN^mCherry), lacking the N-terminal CH domain, failed to localize in the cilia (Fig S2Aa). Also, we noted that the overexpression of either EB1 or EB1^DN^ did not affect the GFPαTub84B enrichment and the cilia volume at 90 hrs APF (Fig. S2Ab-c), indicating that the overexpression of EB1^DN^ was not sufficient to perturb ciliary integrity..

*Drosophila* genome codes for five EB1 (CG3265) homologues (CG32371, CG18190, CG15306, CG2955, Mst27D), of which three (CG3265 >CG18190 >CG32371) bear the highest homology to the mammalian EB1. Therefore, we knocked down EB1 (CG3265) using various EB1^dsRNA^ expression constructs (Fig S3) and found complete loss of *UAS-EB1GFP* expression upon *chaGal4* mediated overexpression of two different *UAS-EB1*^*dsRNAs*^ (BL36599 and BL28605). We used these lines to study the effect on cilia morphology during development and in the adult stage in a tissue-specific manner. The expression of BL28605 by *chaGal4*, lead to significant decrease in the eGFP localization and cilia volume one-day After Eclosion (1 d AE, Fig 2A, and S2B), suggesting that the disruption of EB1 could affect the structural stability of the cilia in the long term.

**Fig 2.**
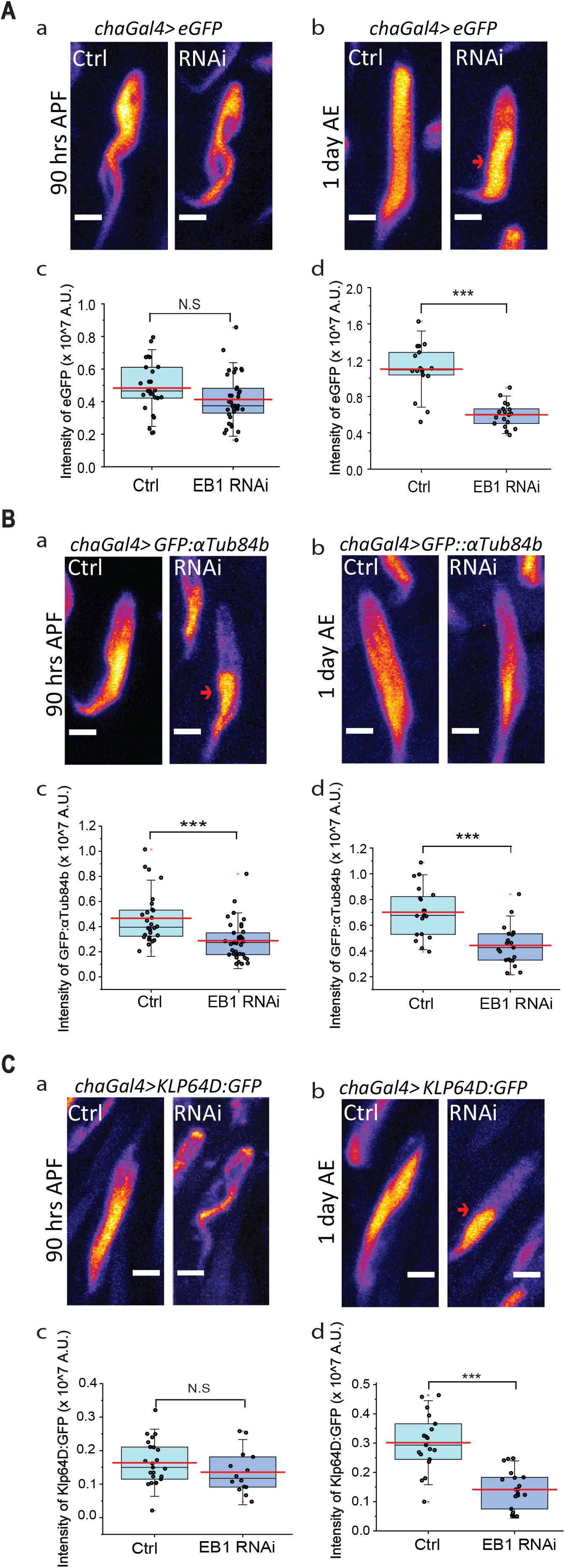
Loss of EB1 in the OSNs disrupts the assembly of the ciliary outer segment and localization of GFPα Tub84B. **A)** Confocal images show eGFP localisation at 90 hrs APF **(Aa)** and 1 day after eclosion **(AE, Ab)**, respectively, in control and EB1RNAi conditions. The intensity of eGFP in the cilia, at 90hr APF **(Ac)** and 1 day AE **(Ad)**, respectively, in the indicated genetic backgrounds. **B)** Confocal images show GFP:αTub84b localisation at 90 hrs APF **(Ba)** and 1 day AE **(Bb)**, respectively, in control and EB1RNAi conditions. The intensity of GFP:αTub84b in the cilia, at 90hr APF **(Bc)** and 1 day AE **(Bd)**, respectively, in the indicated genetic backgrounds. **C)** Klp64:DGFP localisation in the cilia at 90 hrs APF **(Ca)** and 1 day AE **(Cb)**, respectively, in control and EB1RNAi conditions. The Intensity of Klp64D:GFP in the cilia in control and upon EB1RNAi at 90 hrs APF **(Cc)** and 1 day AE **(Cd)**, respectively. The pair-wise test of significance of difference was calculated using Kruskal-Wallis ANOVA, (p<=0.05 *, p<=0.01 **, p<=0.001 ***). The arrows show localization in the OS. Scale bar ∼ 2 μm.

Further, we noted that EB1 RNAi significantly reduced the GFPαTub84B enrichment in the cilia at both 90 hrs APF and 1 d AE (Fig 2B, and S2C). The reduction of GFPαTub84B from the cilia in the absence of EB1 may indicate either a loss of soluble tubulin or microtubule. To distinguish between these two possibilities, we estimated the kinesin-2 localization in the cilia using KLP64DGFP. It decorated the entire length of the cilia in wild-type control at both 90 hrs APF and 1 d AE, suggesting a uniform microtubule distribution along the ciliary OS (Fig 2C-a, b). The EB1 RNAi significantly reduced the KLP64DGFP localization in the cilia at 1 d AE and the motor failed to reach the distal part of the ciliary OS (Fig 2C-d, S2D). KLP64D forms an obligate heterotrimer with KLP68D and DmKAP to make a stable and functional kinesin-2 motor complex and loss of both Klp64D and DmKap affected the cilia growth in *Drosophila* (5,13,32). Therefore, we reasoned that KLP64DGFP could only be localized in the cilia due to its microtubule-binding ability and motor activity. Hence, reduced enrichment of GFPαTub84B and KLP64DGFP in the EB1 RNAi background would suggest that loss of EB1 could affect both the tubulin enrichment and microtubule assembly in the ciliary OS during development and the protein is required to maintain the cilia stability in the adult stage.

### Tubulin and EB1 enters in phase-separated pulses during the assembly of the ciliary outer segment

The kinesin-2-dependent intraflagellar transport (IFT) is implicated in the tubulin entry into the primary cilia and flagella (7,33). Kinesin-2 motors were also shown to directly bind αTubulin1A isoform (which includes αTubulin84B of *Drosophila*) and transport αTubulin84B into the OS of olfactory cilia in *Drosophila* (13). In contrast, studies in *Chlamydomonas* flagella suggested EB1 diffuses into the flagella/cilia and gets enriched at the tip by microtubule capture (9). However, the results discussed above suggest that both tubulin and EB1 localization in the bipartite sensory cilia are developmentally regulated.

Therefore, to understand the underlying mechanisms, we observed the GFPαTub84B and EB1GFP localization patterns in the individual cilium expressed on the Or92A and Or42B OSNs at hourly intervals during 80-90 hrs APF (Fig. 3A-D). The Or92A cilium had two significant GFPαTublin84B surges at 83 hr and 86 hr APF (Fig 3E), whereas a single surge was observed in Or42b cilium at 85hr APF (Fig 3F). Interestingly, the GFPαTub84B surges were always phase-separated by an hour from that of the EB1GFP and the tubulin surge always followed that of the EB1. The phase-separated pulsatile entry of GFPαTubulin84B and EB1GFP indicated that the entry and exit of tubulin and EB1 are independently regulated during the ciliary OS growth in an OSN-specific manner. Also, the pulsatile GFPαTubulin84B surges suggested that the microtubule cytoskeleton of the ciliary OS is dynamic during the development. However, both the spatial and temporal resolutions of the images were inadequate to determine whether the tubulin and EB1 enter the cilia in a particulate form, such as by using the IFT particle, or individually through diffusion. Nevertheless, the phase separation between the tubulin and EB1 pulses raised an intriguing question – whether the EB1 localization promotes entry of tubulin or vice versa.

**Fig 3:**
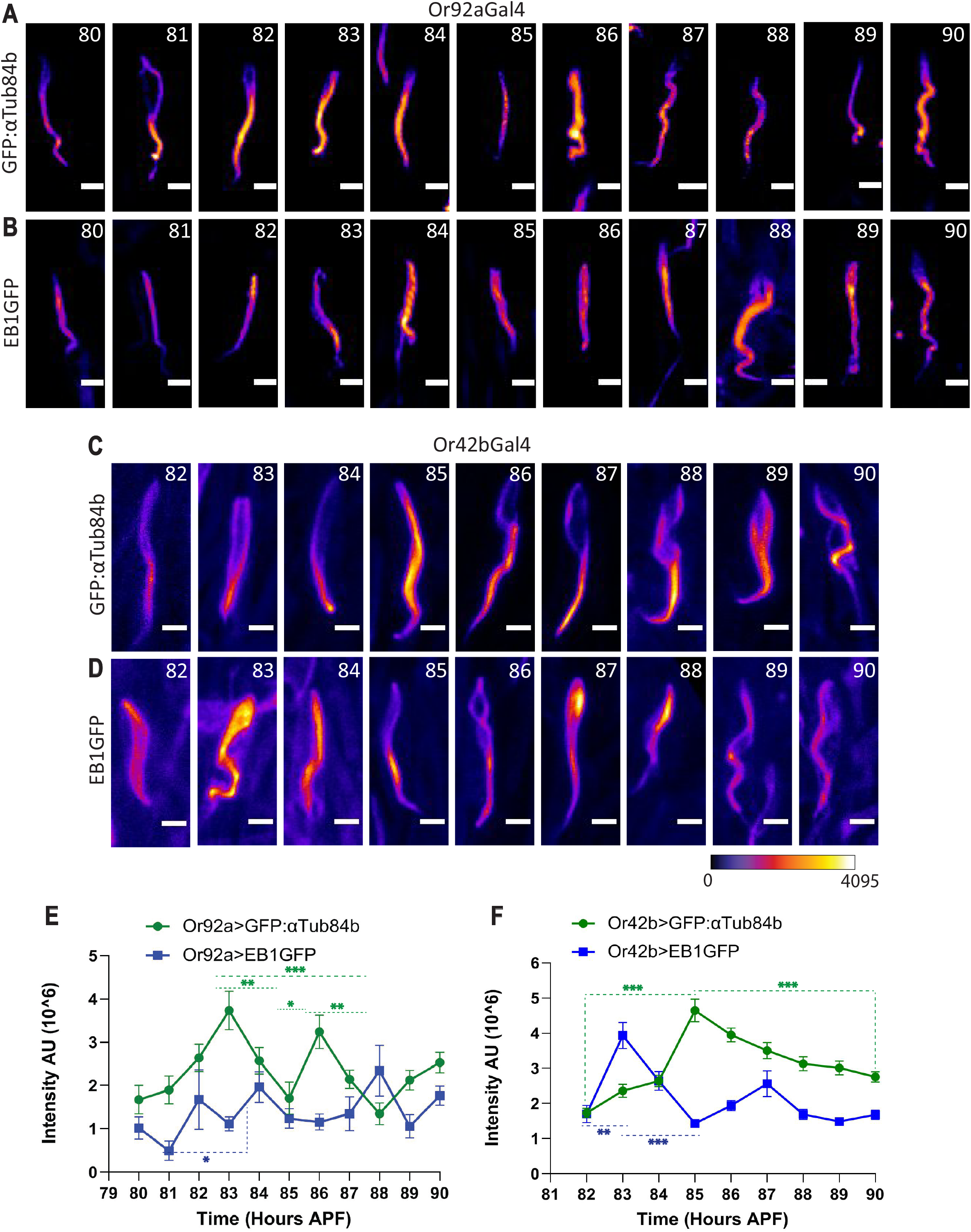
A comparative analysis of the GFPαTub84B and EB1GFP enrichments in the ciliary OS in Or92A and Or42B cilia during development. **A-D)** Representative images of GFPαTub84B and EB1GFP localisations in the Or92a and Or42b cilia during the indicated pupal stages. **E and F)** Integrated fluorescence intensity (mean +/-S.E.M) of GFPαTub84B and EB1GFP localisations in Or92a and Or42b cilia during the 80-90 hr APF. The pairwise significance of difference was estimated using Kruskal-Wallis ANOVA test and the p-values (*p<0.05, **p<0.01, and ***p<0.001) are shown on the plots. Images are shown in false colour intensity heat map (FIRE, ImageJ) and scale bars indicate ∼2 µm each.

### EB1 is required for pulsatile tubulin entry into individual cilium during development

To address the issue, we monitored the GFPαTubulin84B localization dynamics into individual Or92A and Or42B cilia in tissue-specific EB1 RNAi backgrounds during 82-90 hrs APF in cilia (Fig. 4). The EB1^dsRNA^ was expressed at 72 and 64 hrs APF, respectively, in Or92A and Or42B OSNs, using the *Or92aGal4* and *Or42bGal4* drivers (Fig S4A-B), to ensure a late knockdown. This strategy also helped rule out any residual effect due to the loss of EB1 at an earlier stage during the initial growth of the cilia. The loss of EB1 visibly reduced GFPαTubulin84B localisation during 81-85 hrs APF and produced abnormal accumulations during the later stages in the Or92A cilia (Fig. 4A). A similar phenotype was observed in the Or42B cilia (Fig 4B). Intensity quantification showed a total abolition of the GFPαTubulin84B surges during 80-90 hrs APF in both the Or92A and Or42B cilia (Fig. 4C, D), even though the ciliary structure appeared intact. Thus, EB1 is indicated to play a significant role in tubulin entry and/or regulating the microtubule dynamics in the cilia. The tubulin entry is likely to be regulated through association with the IFT particles or independently through motor-mediated transport. Both these processes would require kinesin-2. Hence, we conjectured that the EB1 decoration could stabilize the ciliary microtubule to facilitate further transport along the OS and potentially promote new microtubule assembly.

**Fig 4:**
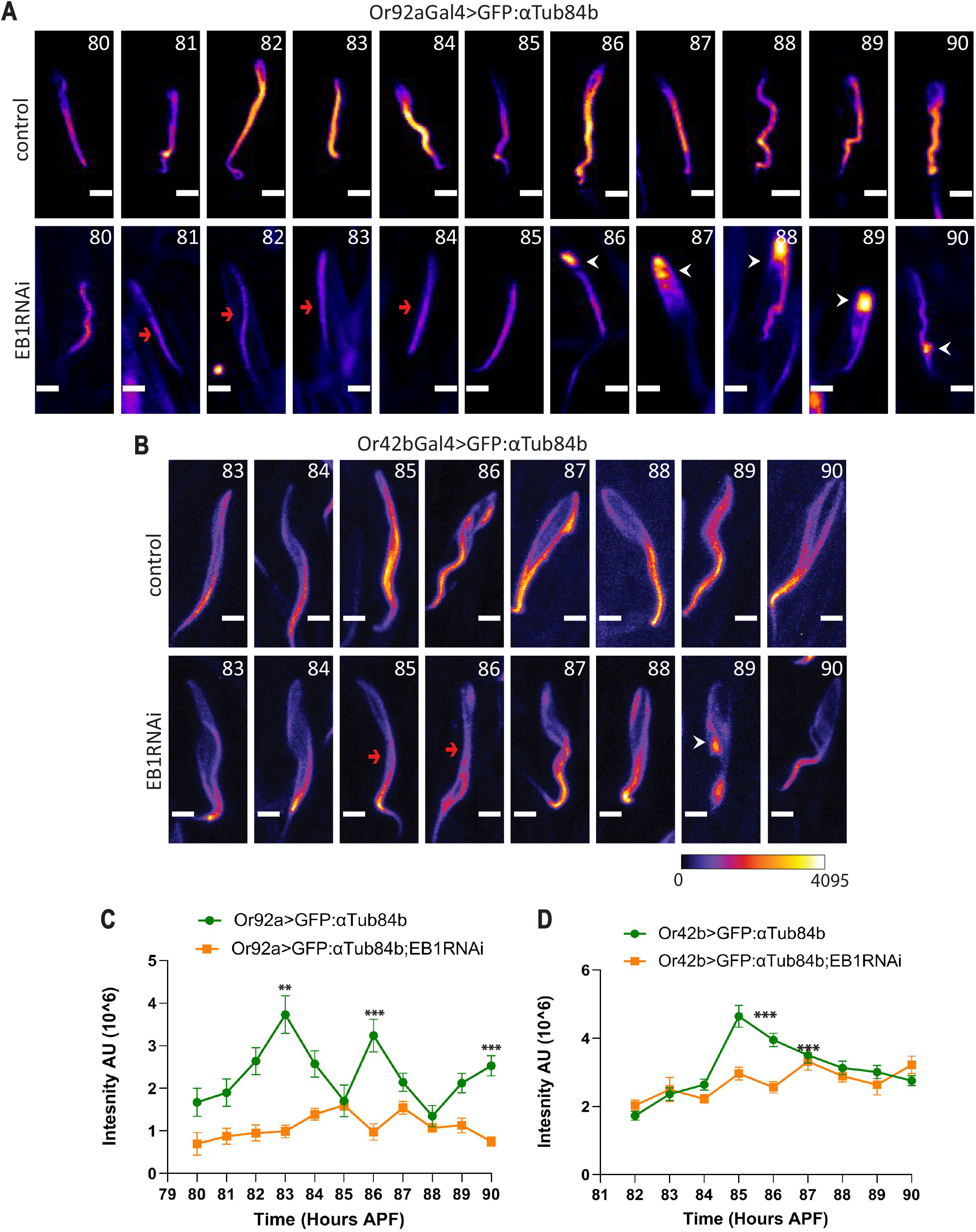
Effects of tissue-specific EB1 RNAi on the GFPαTub84B localization in the ciliary OS. **A and B)** GFPα Tub84B localisation in the ciliary OS of the Or92a (A) and the Or42b (B) cilia in control (top) and EB1 RNAi (bottom) backgrounds during 80-90 hrs APF. Arrows indicate reduction in GFPαTub84B intensity at the indicated developmental time points upon EB1RNAi and arrowheads indicate abnormal accumulation of GFPαTub84B in the RNAi background. **C and D)** GFPαTub84B fluorescence intensity (mean +/-S.E.M) in the ciliary OS of Or92A **(C)** and Or42B **(D)** cilia during 80-90 hrs APF. The pairwise significance of difference was estimated using the Kruskal-Wallis ANOVA test, and the p-values (*p<0.05, **p<0.01, and ***p<0.001) are indicated on the plots. Images are shown in false colour intensity heat map (FIRE, ImageJ) and scale bars indicate ∼2 µm.

### EB1 selectively interacts with KLP68D tail domain of Kinesin-2

Further, the developmentally regulated entry and exit of EB1 indicated that it is actively recruited inside olfactory cilia in response to developmental cues, instead of a free passive diffusion. EB1 associates with its target proteins through the consensus ‘SxIP’ motif found in the tail domain of many partner proteins like Kif17, a kinesin-2 family motor, and the Dynactin P150. Further, EB1 was shown to directly bind to the KIF17 tail domain (34). We found an ‘SSIP’ motif in the tail domain of KLP68D (Fig S5A), which is partly homologous to Kif17, raising a possibility of a direct interaction between the motor and EB1. Affinity copurification using the GST-tagged KLP68D tail fragment (GST-KLP68DT) as baits pulled down EB1GFP from the cell-free extracts of *Drosophila* heads expressing the *UAS-EB1:GFP* transgene (Fig 5A). We used GST-KLP64DT lacking the ‘SxIP’ motif as the negative control. A reverse pulldown experiment using the FLAG-tagged EB1 as bait, which was co-expressed in *Drosophila* neurons along with transgenic KLP68DYFP and the tail-less KLP68DΔTYFP, respectively, only copurified the full-length KLP68DYFP (Fig 5B). Also, EB1GFP failed to bind the human kinesin-2 orthologues GST-KIF3AT and GST-KIF3BT as both the tail domains lacked the ‘SxIP’ motif (Fig S5B). Altogether the data indicated that EB1 can bind to the KLP68D tail both in vitro and in vivo. Thus, it raised the possibility of kinesin-2 could directly transport EB1 into the cilia.

**Fig 5.**
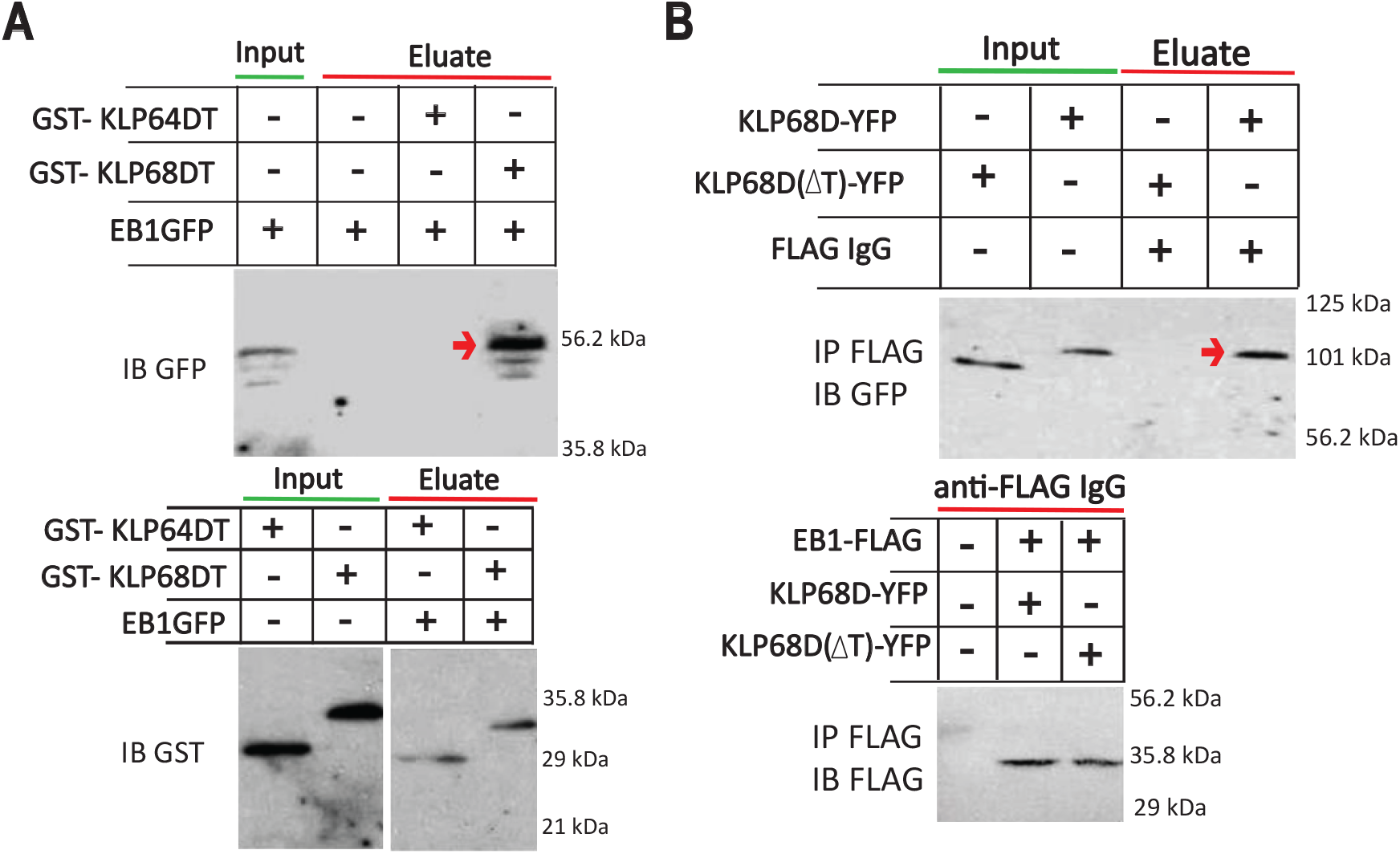
Selective EB1 interaction with the KLP68D tail domain of Drosophila kinesin-2 in vitro. **A)** Copurification of EB1:GFP from the chaGal4>EB1:GFP expressing Drosophila head extracts with the recombinant Glutathione S-transferase (GST)-tagged, KLP68D tail (GST-KLP68DT) fragment using affinity chromatography. The arrow indicates the EB1:GFP band. **B)** The immune-coprecipitation (IP) of the recombinant KLP68D from the head extracts of chaGal4>Klp68D:YFP and chaGal4>Klp68D(ΔT)YFP, expressing UAS-EB1FLAG using anti-FLAG. The arrow indicates a full-length KLP68DYFP band.

### Transient EB1 enrichment in the olfactory cilia requires kinesin-2

To test this possibility, we disrupted kinesin-2 function in Or92A and Or42B OSNs shortly before the 80-90 hrs APF interval by expressing *UAS-Klp68D*^*dsRNA*^ (GD-27944) in these neurons. Although the total EB1GFP localisation was not severely affected upon the Klp68D RNAi in both the Or92A and Or42B cilia during the growth interval (Fig. 6A and B), we found several abnormal EB1GFP accumulations along the length of the cilia at every developmental stage. Also, the EB1GFP surges at both the 84 hrs and 88 hrs APF were significantly reduced in the Or92A cilia upon Klp68D RNAi (Fig 6C). A similar reduction was observed in the Or42B cilia in the Klp68D RNAi background at 83 hrs APF (Fig 6D). Together these data suggested that kinesin-2 is required for the EB1 entry into the cilia. The abnormal accumulation observed along the OS length may further indicate an involvement of kinesin-2 in maintaining the EB1 distribution. Alternatively, it could be attributed to the apparent loss of microtubule along the ciliary OS. This result is in apparent contrast to the earlier report which suggested that EB1 could diffuse along the *Chlamydomonas* flagella (9). Here it would be important to note that unlike the olfactory cilia, which contains a bipartite microtubule architecture with a 9+0 doublet-microtubule bearing CC and singlet microtubule-bearing IS and OS, the flagella and motile cilia contain only doublet microtubule-bearing 9+2 axoneme. EB1 enriched only at the distal tips of the flagella and motile cilia, whereas it decorated the singlet microtubule-bearing distal parts of the MDCK cilia and the OS of olfactory cilia (21,25). Further interaction between KIf17 is demonstrated to localize EB1 at the growing tips of cellular microtubules and stabilize them (34). Therefore, the pulsatile increments of EB1 along the OS could periodically stabilise the singlet microtubules, and its active outflux could promote microtubule dynamics. This hypothesis is consistent with the observed loss of KLP64DGFP from the distal OS in the *cha>EB1* RNAi background (Fig 2). Also, Kinesin-2 could play a significant role in the process by inducing periodic influx of EB1 into the cilium.

**Fig 6:**
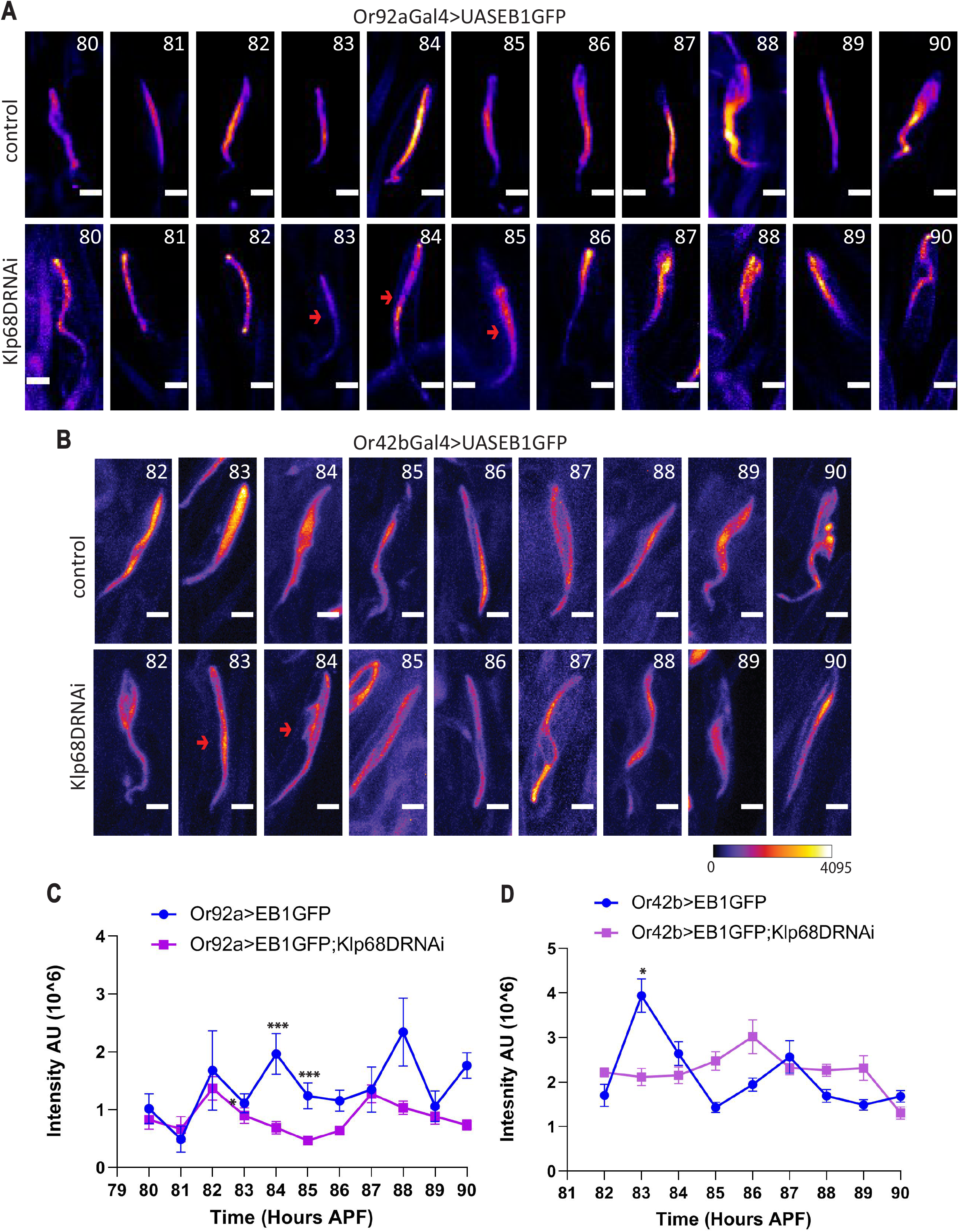
Effects of tissue-specific Klp68D RNAi on the EB1GFP localisation in the ciliary OS. **A and B)** EB1GFP localisation in the ciliary OS of the Or92a (A) and the Or42b **(B)** cilia in control (top) and Klp68D RNAi (bottom) backgrounds during 80-90 hrs APF. Arrows indicate loss in EB1GFP intensity in the RNAi background at the indicated developmental timepoints. **C and D)** EB1GFP fluorescence intensity (mean +/-S.E.M) in the ciliary OS of Or92A **(C)** and Or42B **(D)** cilia during 80-90 hrs APF. The pairwise significance of difference was estimated using Kruskal-Wallis ANOVA test, and the p-values (*p<0.05, **p<0.01, and ***p<0.001) are indicated on the plots. Images are shown in false colour intensity heat map (FIRE, ImageJ) and scale bars indicate ∼2 µm.

### Kinesin-2 is required for the tubulin surges during 80-90 hrs APF in the olfactory cilia

To test these possibilities, we monitored the developmental localization of GFPαTubulin84B in the Or92a and Or42b cilia in the Klp68D RNAi backgrounds. The GFPαTubulin84B localisation was visibly reduced, particularly along the distal parts of the Or92A and Or42B ciliary OS in the tissue-specific RNAi backgrounds (Fig.7A and 7B). Levels were significantly lower at 82-83 and 86-87 hrs APF in Or92A cilia, and at 85-86 hrs APF in the Or42B cilia (Fig. 7C, D). We also investigated the effects of tissue-specific, Klp68D and EB1, RNAis on GFPαTubulin84B localisation in the Or42b and Or92a cilia in the adult stage (1hr AE) (Fig S6). It revealed a significant loss of tubulin enrichment in the respective ciliary OSs (Fig S6), indicating that both EB1 and KLP68D are essential for the tubulin accumulation in the ciliary OS. Interestingly, in both these cases the tubulin enrichment was affected at the distal part of the OS (Fig S6A, B). The result, therefore, would suggest that kinesin-2 function is necessary for tubulin entry and distribution along the length of the ciliary OS. Overall, the data is consistent with the findings that tubulin movement inside the cilia/flagella requires the IFT and kinesin-2. However, it was unclear whether the tubulin surges in the cilia would indicate transient microtubule growth.

**Fig 7:**
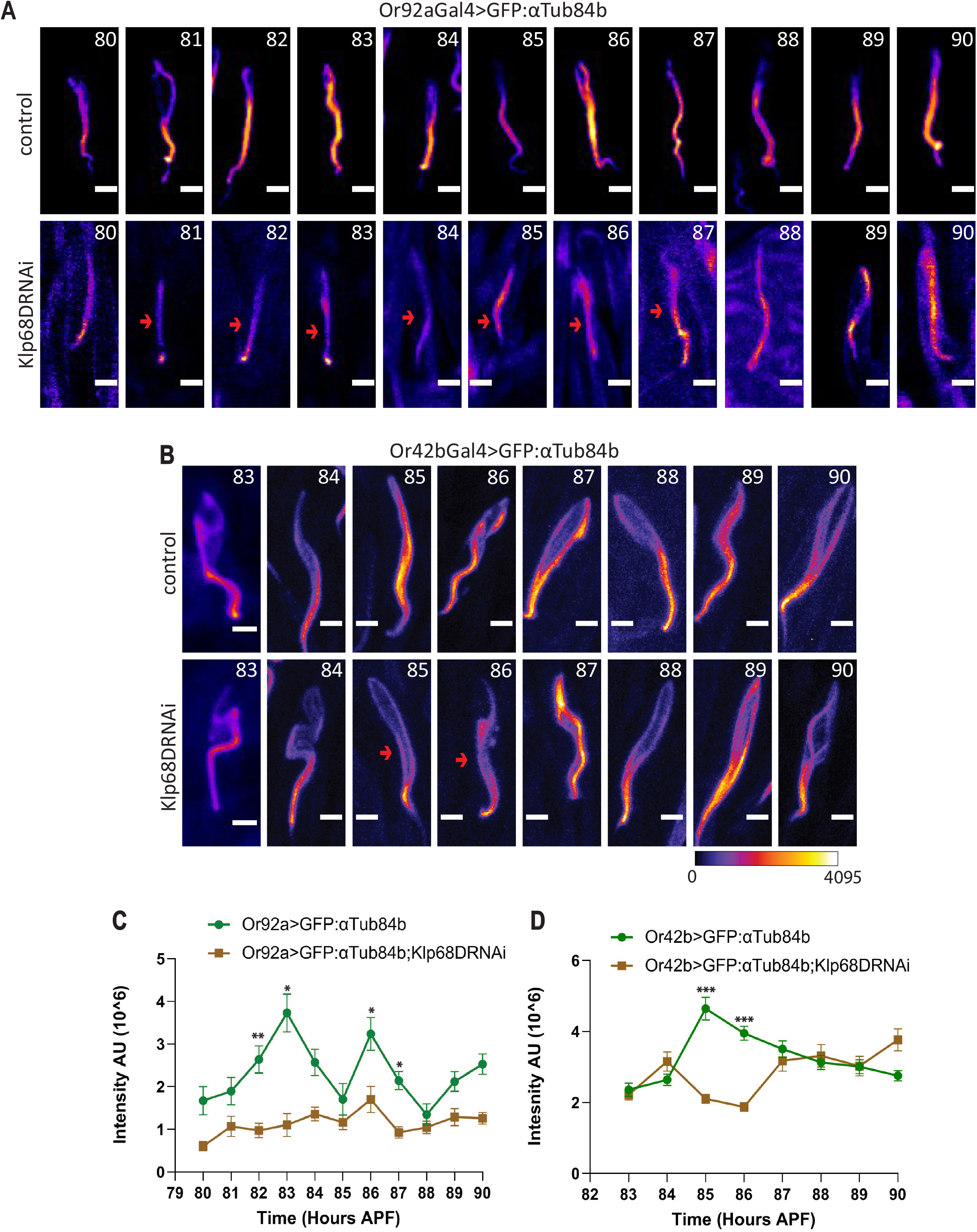
Effects of tissue-specific Klp68D RNAi on the GFPαTub84B localisation in the ciliary OS. **A and B**) GFP αTub84B localisation in the ciliary OS of the Or92a **(A)** and the Or42b **(B)** cilia in control (top) and Klp68D RNAi (bottom) backgrounds during 80-90 hrs APF. Arrows indicate loss of GFPαTub84B localisation in the ciliary OS upon Klp68D RNAi at the indicated timepoints. **C and D)** GFPαTub84B fluorescence intensity (mean +/-S.E.M) in the ciliary OS of Or92A **(C)** and Or42B **(D)** cilia during 80-90 hrs APF. The pairwise significance of difference was estimated using Kruskal-Wallis ANOVA test, and the p-values (*p<0.05, **p<0.01, and ***p<0.001) are indicated on the plots. Images are shown in false colour intensity heat map (FIRE, ImageJ) and scale bars indicate ∼2 µm.

**Fig 8.**
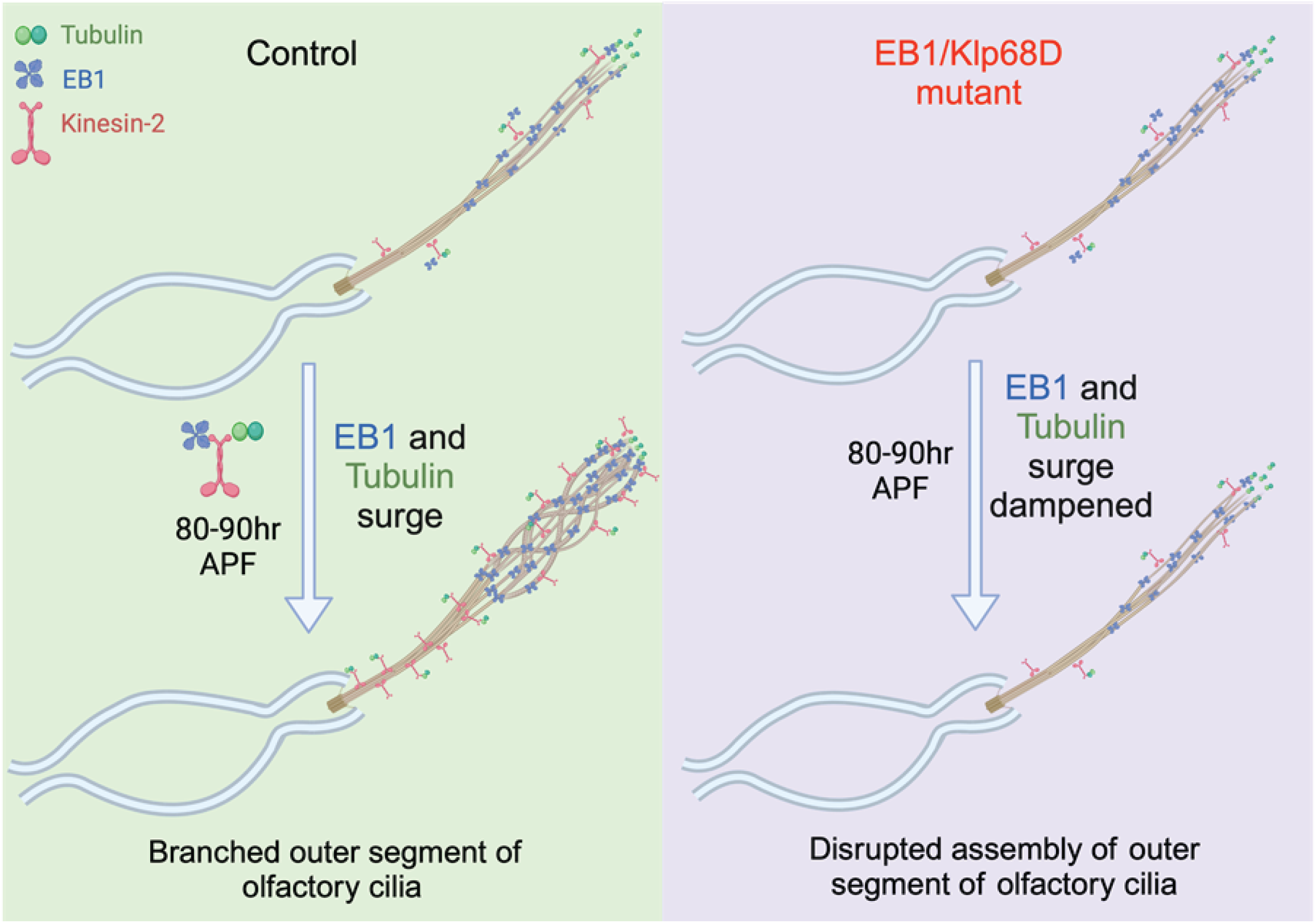
Schematic illustrates the EB1 and tubulin transport mechanisms in the growing olfactory cilia in the wild type and mutant backgrounds. The data presented in previous figures and by Girotra et al., (2017) suggest that EB1 and Tubulin are transported by the kinesin-2 motor into cilia through a possible direct interaction between KLP68D and EB1 (shown here) and tubulin (Girotra et al., 2017) during the outer segment (OS) growth in each type of bipartite olfactory cilia. Loss of EB1 transport affects tubulin localisation and OS growth and stability.

## Discussion

The ciliary axoneme is made up of doublet microtubules arranged in a ninefold circular symmetry represented as 9+0 or 9+2 arrangements (2). In most cell types the cilia or flagella grow in a single dimension without showing much diversity in morphology and ultrastructure (1,7). However complex sensory cilia are bipartite and do not retain doublet arrangement to the tip of cilia (4). They have complex architectures in the outer segment supported by singlet microtubules (3). We observed that each olfactory cilia grows independently with distinct growth profiles and developmentally regulated, out of phase localisations of EB1 and tubulin. The neuron-type-specific surges in EB1 and tubulin might influence the morphology of bipartite cilia. A similar observation was made by Howell and Hobert (2017) wherein, the dosage of a transmembrane proteins expressed at different levels in different olfactory neurons was involved in determining extent of cilia elaborations of ciliated sensory endings of *C. elegans* olfactory neurons (35).

Here, we show that pulsatile EB1 entry preceding the tubulin entry and overall growth of the ciliary OS could facilitate the tubulin entry and cilia growth. We also showed that the Kinesin-2 motor could play a significant role in the process through a potentially direct interaction with EB1. In a previous study we had shown that Kinesin-2 could directly interact with αTubulin1A/D and its orthologue, the αTubulin84B of *Drosophila*, in vitro (13). Therefore, we concluded that EB1 entry is necessary to facilitate that of tubulin and subsequent ciliary growth. Inexplicably, we noted certain differences between the ensemble data obtained using *chaGal4>GFP:*_α_*Tub84B and Jupitar-GFP* probes and that obtained using the *Or92aGal4>GFP:*_α_*Tub84B* and *Or42bGal4>GFP:*_α_*Tub84B*. In the first instance the tubulin levels appreciated in steps and retained the level until the next episode. Whereas the latter depicted pulsatile influx of tubulin leading to a volume growth towards the end. It is important to note that *Or92aGal4>GFP* and *Or42bGal4>GFP* probes indicate a net OS volume growths occur during 89-90 and 87-89 hrs APF intervals, respectively, in these two cases, which coincided with the 89-90 hrs APF growth window of the composite cilia. Also, the composite ensemble data includes Or10a and Gr22a (36) cilia which are not measured individually due to lack of adequate reagents. Hence, it would not be feasible to comment on this discrepancy. In spite of this lacunae, the EB1GFP pulses appeared consistently in all three cases. Also, the EB1 surges subsided preceding the tubulin surges in all three reporter systems. Therefore, we decided to interpret the utility of the EB1 surges in the context of tubulin entry and cilia growth.

It is well known that the disruption of EB1 microtubule association could affect the structural stability of the cilia in the long term (37). In *Drosophila* olfactory cilia, the EB1 RNAi significantly reduced the enrichments of GFPαTubulin84B and KLP64DGFP in the ensemble of cilia, indicating loss of EB1 could disrupt microtubule assembly and tubulin enrichment in the cilia. We also observed that even EB1 despite being a small molecular protein (30 kDa) is transported actively in response to developmental cues inside the olfactory cilia. This is unlike the prevailing knowledge in the field where it has been shown that, ciliary base acts like a molecular sieve allowing proteins below 40kDa to diffuse passively inside primary cilia (38). Moreover, EB1 has also been reported to simply diffuse to the tip of the Chlamydomonas flagella during regeneration and it accumulates at the distal tip by a diffusion-and-capture mechanism (9). Since the Chlamydomonas flagella is a simple rod-shaped structure, diffusion-and-capture might suffice to localise small proteins to the tip of the flagella but due to the increased architectural complexity in the ultrastructure of the flagellum in *Drosophila* olfactory cilia, even small proteins like EB1 might have to be transported actively indicating that diverse cilia would require distinctive mechanisms for assembly and maintenance.

The EB1 entry inside olfactory cilia is likely to be an active, motor-mediated process, which may take place either through direct interaction of EB1 with kinesin-2, probably through the ‘SSIP’ motif present in the tail domain, or by association with IFT particles. The KLP68D-dependent EB1 localisation along the OS could stabilise the singlet microtubules and its loss could promote dynamics as loss of KLP68D leading to the abrogation of EB1GFP surges and abnormal accumulation of GFPαTubulin84B, a *bona fide* Kinesin-2 cargo, in many other ciliary systems (13,39). The IFT-dependent transport of Tubulin was upregulated during flagellar assembly in regenerating Chlamydomonas flagella. Some of the Tubulin could enter the flagella through diffusion (7). Tubulin directly interacts with the IFT complex through the Tubulin binding module formed by IFT74 and IFT81 subunits during cilia assembly (40). In the *C. elegans* amphid cilia, specific Tubulin isotypes moved along with the IFT particles indicating that IFT transports these Tubulin isotypes to ciliary tips (41). Altogether, these results suggested that Tubulin enters the cilium through two modes-diffusion and active transport by IFT, highlighting that the assembly of axoneme of a complex cilium is an active process.

Several Mitogen-Activated Kinase (MAK) activated by neurotransmitters, cytokines, hormones and growth factors, which transduce developmental cues (42–44)), elongates cilia to varying degrees in *Chlamydomonas, C. elegans*, and mammalian cells (45–48). Disrupted kinase function abolishes length-dependent modulation of tubulin transport in *Chlamydomonas* flagella (7,49). In mouse photoreceptors, MAK has been reported to regulate MT stability and the length of connecting cilia (48), in AWA cilia of *C. elegans* DYF-18/CCRK and DYF-5/MAK regulate axoneme length and ciliary branches (50). However, the targets of the kinases that regulate ciliary architecture are not known. Mutations of dyf-5 and dyf-18 showed ectopic localization of the kinesin-2 at the ciliary distal segments (51), and DYF-5 mutation affects docking and undocking of kinesin-2 motors and reduces their speed in the cilia of *C. elegans* (47). The MAK family kinases have also been shown to directly phosphorylate kinesin-2 and the RP1 MT-binding protein (48,52). Hence the kinases might regulate the axoneme length and arborisation by regulating the kinesin-2 localisation, its interaction with IFT particles, and docking on the ciliary axoneme or the velocity of the motor. Therefore, the kinesin-2 mediated surges of EB1 and tubulin in *Drosophila* olfactory cilia could be developmentally regulated by kinases. Further, some kinases like ICK and MOK kinases have been reported to regulate the levels of soluble tubulin via modulation of mTORC1 activity in renal epithelial cells (52–54). Thus, these developmentally regulated kinases may target different proteins and mechanisms to tune the length of diverse cilia across species. Continued investigations can provide more insight into the molecular mechanisms that generate and maintain cilia diversity to enable different cells to perform diverse functions.

## Materials and Methods

### Drosophila culture and stocks

Fly stocks used in the study are mentioned in (Table S1). Flies were reared at 25°C for all the experiments on standard cornmeal agar. The experiments were performed on dissected pupal antennae of different ages according to the experimental requirements.

### Drosophila sample preparation

For live imaging of *Drosophila* pupae, antennae were dissected and mounted in a drop of grade 700 Halocarbon oil (Sigma Chemical Co., MO, USA). For imaging of aged pupal antennae, the wandering-stage third-instar larvae were monitored every 30 minutes until they became stationary with the head and mouth hook stopped moving. The time point was marked as 0 hours After Pupa Formation (0 h APF) stage. The pupal case was opened, and the second and third segments of the antennae were pinched off and fixed in 200 µl of fixative solution (4% Paraformaldehyde in Phosphate Buffered Saline, pH 7.4) for 20 minutes at room temperature, followed by three washes with Phosphate Buffered Saline, pH 7.4 (PBS). Antennae were then mounted in a drop of Vectashield® (Vector Laboratories Inc., USA) on a glass slide under a 0.17 mm coverslip

### Image acquisition and analysis

All fluorescence images were obtained under constant acquisition conditions in Olympus (Olympus Imaging Corp., Japan) confocal microscope FV1200 using a 60x oil 1.4 NA objective lens or FV3000 using a 60x oil 1.4 NA objective lens. Subsequently, all images were processed using ImageJ (rsweb.nih.gov/1 IJ) and presented using the false-colour heat map. Images placed within a figure panel and used as data sets for statistical comparisons between different developmental stages and genetic backgrounds were collected using identical laser power and gain settings to maintain comparability. All Figures containing images and line arts were composed in Illustrator (Adobe Inc., USA).

### Protein extraction from Drosophila heads

Fly heads were manually separated using a razor blade and homogenised in 2 μl per head ice-cold lysis buffer. Each sample, consisting of 150-200 heads, was homogenised using a motorised tissue grinder (Genetix, Sigma Chemical Co. MO, USA). The homogenate was centrifuged at 1,00,000 x g for 60 minutes at 4°C. The supernatant fraction was collected and used in further experiments as soluble, cell-free extracts.

### Pulldown

Cell-free extracts from *Drosophila* heads and recombinant proteins purified through affinity chromatography were mixed appropriately in a wash buffer (TrisHCl – 20mM, MgCl2 – 5mM, NaCl – 150mM, DTT – 1mM) as the reaction mixture. This mixture was incubated at 23°C for 1 hour on a rotor set at 10 rpm. This was then mixed with either previously equilibrated Ni-NTA Agarose beads (Qiagen Ltd, USA) or Glutathione Sepharose beads (GE Healthcare Ltd) in accordance with the tag on the recombinant bait proteins of interest, bead for 1-hour at 23°C at 10 rpm. This suspension was then centrifuged at 15000 rpm and the supernatant was discarded. The bead pellet with bound proteins was washed 3 times with the same rection buffer to reduce nonspecific binding and then eluted with appropriate elution buffers. The eluates were mixed in equal proportions with 2x Laemmli Buffer, boiled, and analysed using SDS-PAGE, Western Blotting and immunostaining.

### Immunoprecipitation

Cell-free extracts from *Drosophila* heads were incubated with a 50% slurry of equilibrated Protein A-Sepharose beads (GE Healthcare Ltd) in the ratio of 100 µl bead slurry for 1 ml extract for the Pre –clearing step in Immunoprecipitation. This mixture was incubated at 4°C for 1 hour with gentle rocking and then centrifuged at 12000 g at 4°C for 20 seconds. The supernatant was collected and incubated with 1–5 µg of appropriate primary antibody and incubated for 1 hour at 4°C with gentle rocking for the Antigen-Antibody coupling stage of Immunoprecipitation. This suspension was mixed with previously equilibrated Protein A-Sepharose beads and again set for 1-hour incubation at 4°C with gentle rocking. This suspension was centrifuged at 12000 g at 4°C for 20 seconds for precipitation of bound Immune Complex. The pelleted beads were resuspended in equal proportions with 2x Laemmli Buffer, boiled, and analysed using SDS-PAGE, Western Blotting and immunostaining.

### SDS PAGE and western blotting

Samples boiled in 2x Laemmli Buffer was run on a 10% Acrylamide gel. Proteins from these gels were transferred onto a previously activated PVDF membrane (Hybond-P, GE Healthcare Ltd) in an electro-blotting apparatus (Bio-Rad, USA) following the supplier’s protocol and incubated in different primary antisera solutions as the following: anti-GFP Rabbit (dilution, 1:500; #3999 Bio Vision Inc., CA, USA) or anti-FLAG Rabbit (1:500; #F7425 Sigma-Aldrich, USA) Or anti-GST mouse (1:1000, Bioklone Biotech Pvt. Ltd.) in 20 mM Tris-buffered saline (TBS, pH 7.4) containing 0.1% TweenR 20. Subsequently, they were incubated either Goat anti-Rabbit:HRP (1:10,000, Bangalore Genei Pvt. Ltd, India), or Goat anti-mouse:HRP (1:20000 dilution, Bangalore Genei Pvt. Ltd, India), in the TBS-T, and developed by using ECLR chemiluminescence detection kit (GE Healthcare Ltd. USA).

### Statistical analysis

All statistical comparisons were carried out using either student’s T-test, Mann-WhitneyU or Kruskal Wallis one-way ANOVA with p values calculated according to Dunn’s multiple comparisons Test in Graphad Prism9 or Origin. P values for data sets in each figure are indicated in the respective figure legends.

