## Supplementary figure for "Active EB1 surges promote tubulin influx into the growing outer segments of the bipartite olfactory cilia in *Drosophila*"

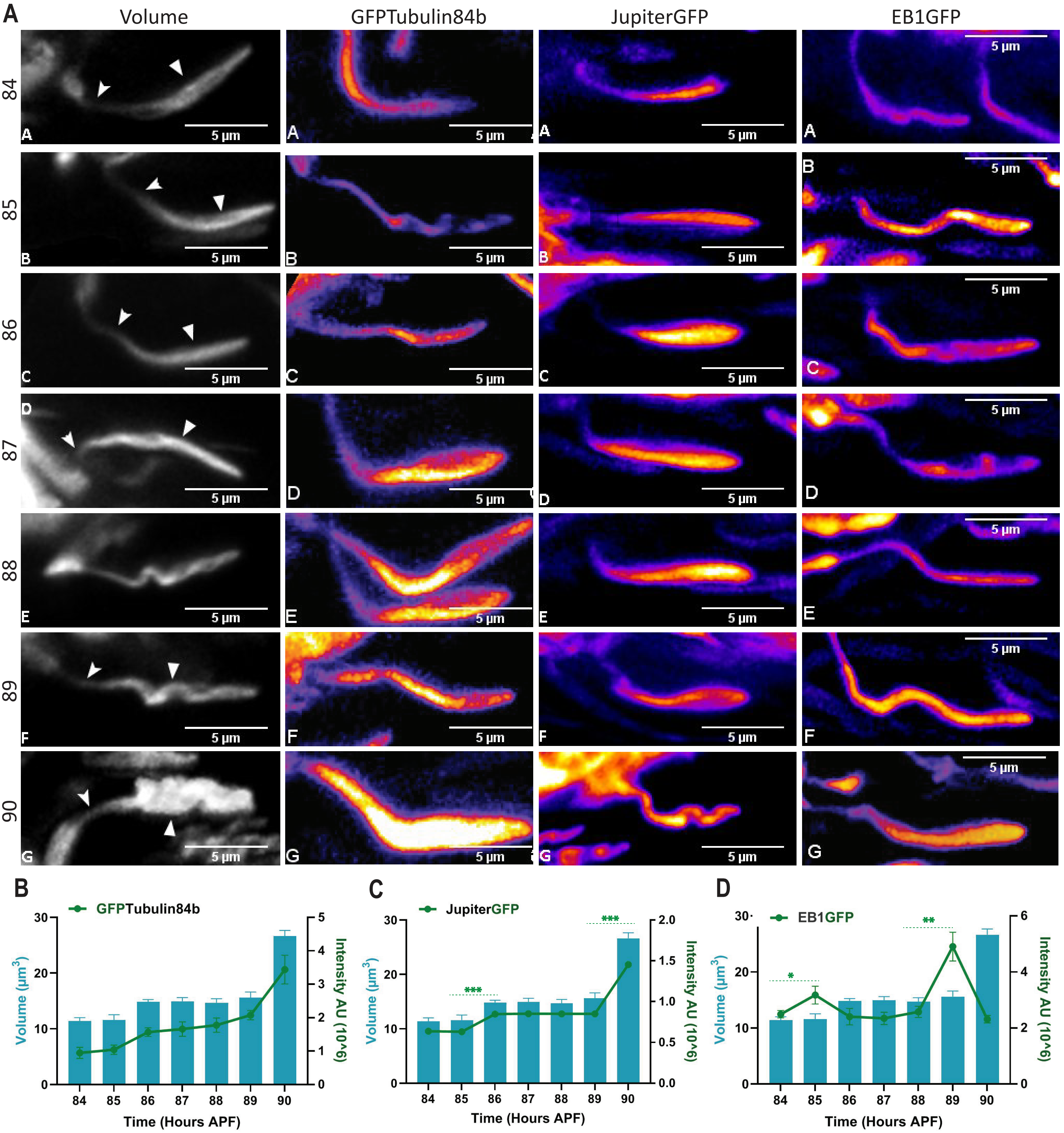


**FigS1: Phasic Growth of cilia inside a basiconic sensilla shaft. A)** Increment of Volume and enrichments of GFPαtubulin84B, JupiterGFP and EB1GFP in AB1-type basiconic cilia at the indicated stages after pupa formation. Notched arrow marks the CC, filled arrow marks the OS. Images are shown in gray scale and false colour intensity heat map (FIRE, ImageJ). **B-D)** Total fluorescence intensities (mean± SEM) of (B) GFPTubulin84b, **(C)** JupiterGFP, and **(D)** EB1GFP in bassiconic cilia expressed using chaGal4 and quantified over a developmental period of 84-90 hr APF. The pairwise significance of difference was estimated using Kruskal-Wallis ANOVA test and the p-values (*p<0.05, **p<0.01, and ***p<0.001) are shown on the plots. Error bars indicate +/- S.E.M.

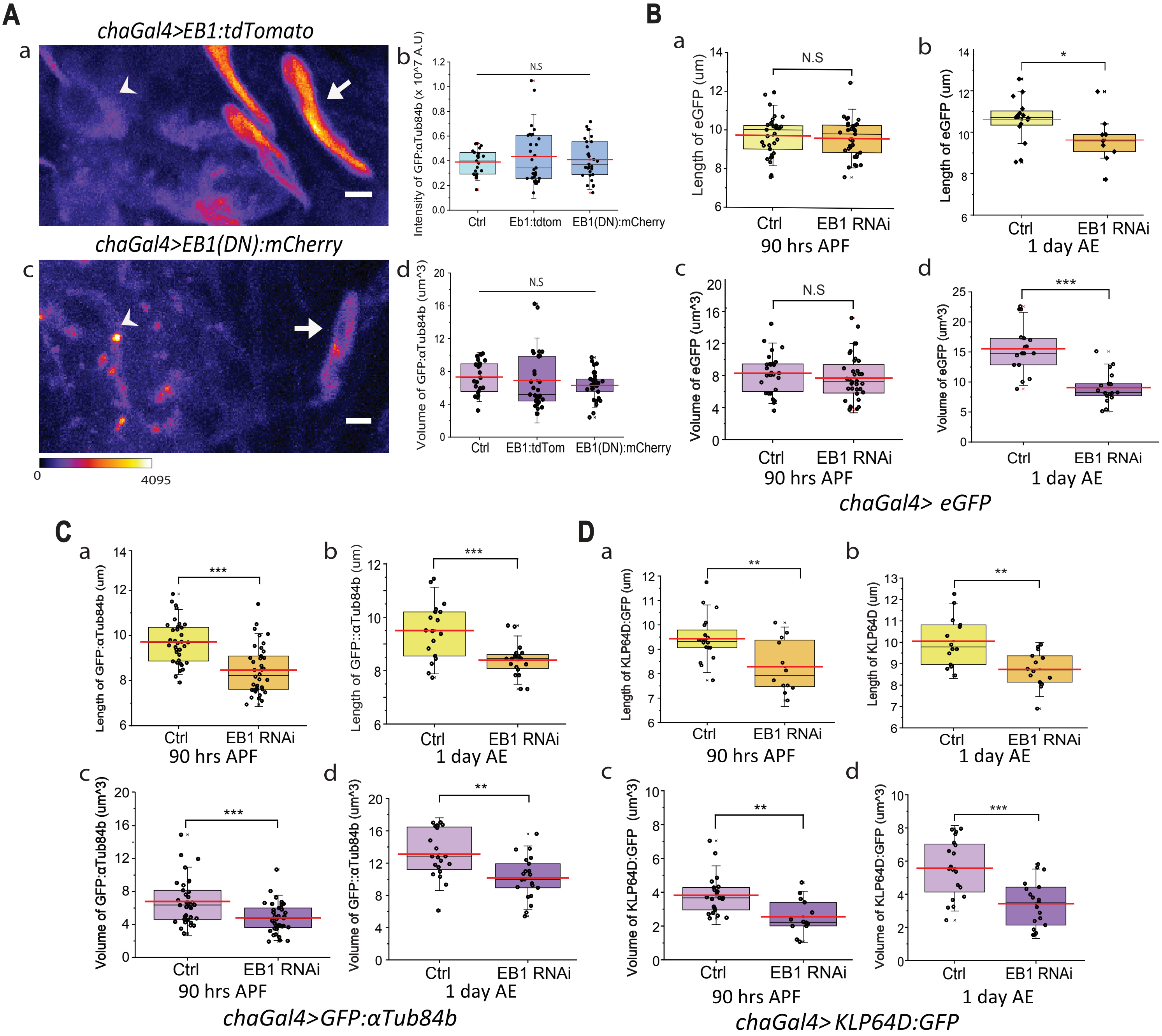


**FigS2: Effect of EB1 RNAi on cilia inside a basiconic sensilla shaft. A)** Localization of GFPαTub84b in the control and the Dominant Negative (DN) backgrounds (a and b). Integrated Intensity (c) and volume (d) of OS marked by GFPα Tub84b, respectively, in control, EB1 overexpression (EB1tdtom) and EB1DN mutants**. B)** length of OS marked by eGFP **(a)** and Volume of eGFP (b), along the length in the ciliary OS in the indicated genetic backgrounds. **C)** Length of outer segment marked by GFPαTub84b (a and b) and Volume of GFPαTub84b marked ciliary OS (c and d) at 90hrs APF and 1day AE. **D)** Length of outer segment marked by KLP64DGFP and (c and d) Volume of KLP64DGFP intensity along the length in the ciliary OS. The pair-wise test of significance of difference was calculated using Kruskal-Wallis ANOVA. p-values (*p<0.05, **p<0.01, and ***p<0.001) are shown on the plots. Error bars indicate + S. D.

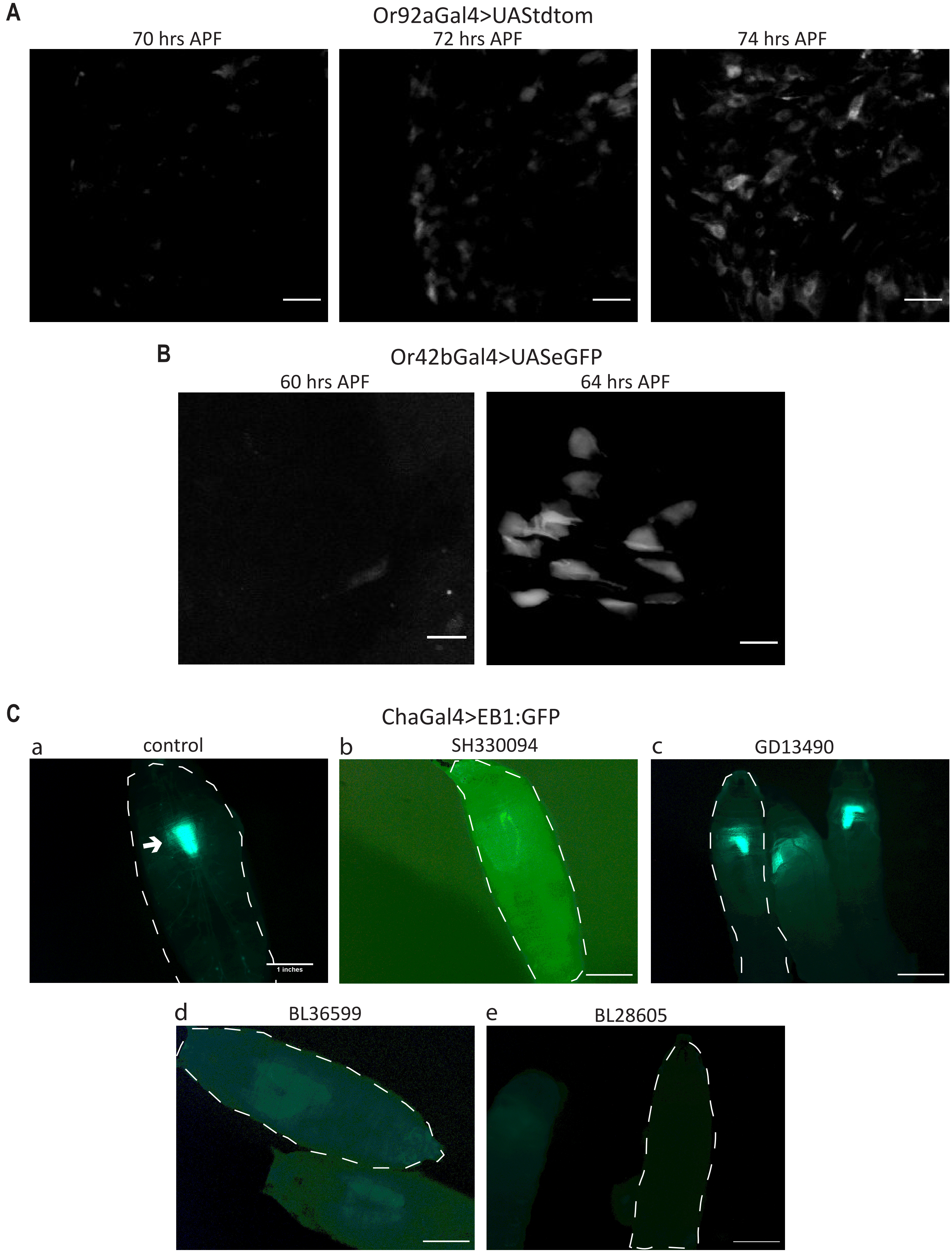


**Fig S3: Validation of the efficacy of the UAS-EB1 dsRNA constructs. A and B**) Or92aGal4 and Or42bGal4 start expressing at 72 hr and 64 hr APF, respectively. The confocal images show Or92aGal4>UAS-tdtom and Or42bGal4>UAS-eGFP expressions at the indicated time points. **(Ca)** EB1GFP expression in the control (cha>UAS-EB1:GFP/ UAS-Dicer). Three different dsRNA stocks - **(Cb)** SH 330094, **(Cc)** GD 13490, **(Cd)**BL-36599 and **(Ce)**BL26805 were tested in the larval stages. The dashed white line shows the outline of the larva and the arrow indicates towards the EB1GFP expression in the ventral ganglion in the control larva. The fluorescence is abrogated in the EB1dsRNA (BL36599 and BL28605) backgrounds.

**A**

| | |
|---|---|
| Hs KIF3A | 600 QDYQEMIENYVHWNEDIGEWQLKCVAYTGNNMRKQTPVPDKKEKDPFEVDLSHVYLAYTEESLRQSLMKLERPRTSKGKARPKTGRRKRSAKPETVIDSLLQ 701 |
| Hs KIF3B | 593 LEEKSKIMNRAFFDEEEDHWKLHPITRLENQQMMKRPVSAVGYKRPLSQHARMSMMIRPEARYRAENIVLLELDMPSRTTRDYEGPAIAPKVQAALDAALQDED EIQVDASSFESTANKKSKARPKSGRKSGSSSSSSGTPASQLYPQSRGLVPK 747 |
| Dm Klp64D | 589 PKEYQSMINQYTHWNEDIGEWQLKCVAYTGNNMRKHISAHKTSGKEPDFLDLSHVYLSYNTDGVSNPMRSKSARPRTSGVPRPTTARRY 677 |
| Dm Klp68D | 584 PKVSASLQAVLAQAMQTGGDDIDIVDSHTNSLRSRLENIINANANGGAGPGAGVAVG**SSIP**NVRNIKSSRGLPSAASNLDSNRRPPTGRLPAKKPASAYPKARGLVNK 784 |

**B**

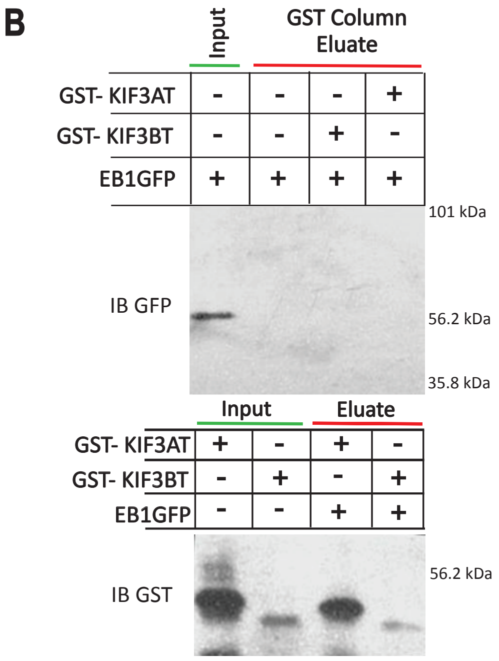


**Fig S4: Pulldown of GFP tagged EB1 by recombinant mammalian GST tagged Kinesin–2 tail fragments. A)** Tail sequences of the motor subunits of Homo sapiens kinesin-2 (HsKIF3A and HsKIF3B) and Drosophila melanogaster kinesin-2 (DmKlp64D and DmKlp68D). A consensus EB1-binding motif (SXIP) in DmKlp68D tail is indicated in red. **(B)** Affinity purification of EB1:GFP from chaGal4>EB1:GFP Drosophila head extract using GST-KIF3AT and GST-KIF3BT.

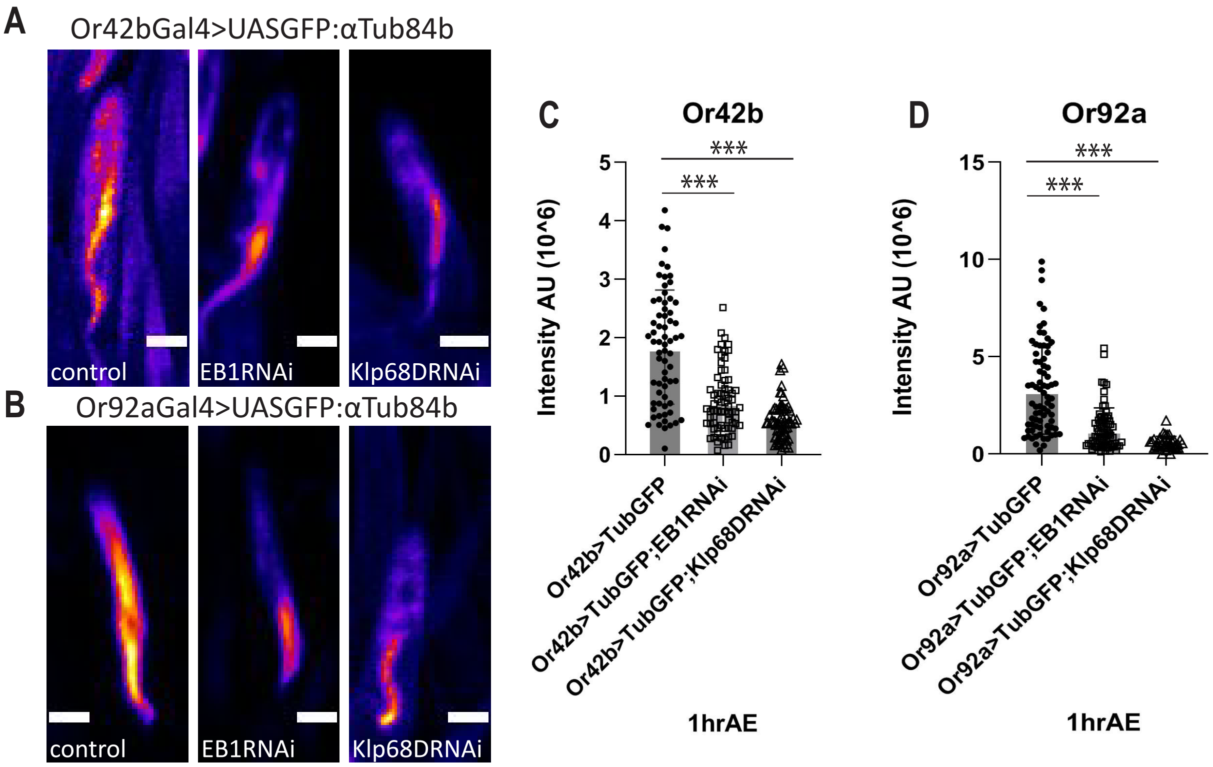


**Fig S5: Effects of EB1 and Klp68D RNAi, respectively, on the tubulin levels in the cilia of adult flies. A and B)** Representative images of GFPαTub84b localisation in Or42b **(A)** and Or92a **(B)** cilia in the indicated genetic backgrounds. **C and D)** Integrated fluorescence intensity (mean +/- S.E.M) of GFPαTub84b localisation in Or42b **(C)** and Or92a **(D)** cilia respectively. The pairwise significance of difference was estimated using unpaired Student's T-test, p-values (*p<0.05, **p<0.01, and ***p<0.001) are shown on the plots. Images are shown in false colour intensity heat map (FIRE, ImageJ). Scale bar indicates 2μm.
